# Disruption in structural-functional network repertoire and time-resolved subcortical-frontoparietal connectivity in disorders of consciousness

**DOI:** 10.1101/2021.12.10.472068

**Authors:** Rajanikant Panda, Aurore Thibaut, Ane Lopez-Gonzalez, Anira Escrichs, Mohamed Ali Bahri, Arjan Hillebrand, Gustavo Deco, Steven Laureys, Olivia Gosseries, Jitka Annen, Prejaas Tewarie

## Abstract

Understanding recovery of consciousness and elucidating its underlying mechanism is believed to be crucial in the field of basic neuroscience and medicine. Ideas such as the global neuronal workspace and the mesocircuit theory hypothesize that failure of recovery in conscious states coincide with loss of connectivity between subcortical and frontoparietal areas, a loss of the repertoire of functional networks states and metastable brain activation. We adopted a time-resolved functional connectivity framework to explore these ideas and assessed the repertoire of functional network states as a potential marker of consciousness and its potential ability to tell apart patients in the unresponsive wakefulness syndrome (UWS) and minimally conscious state (MCS). In addition, prediction of these functional network states by underlying hidden spatial patterns in the anatomical network, i.e. so-called eigenmodes, were supplemented as potential markers. By analysing time-resolved functional connectivity from fMRI data, we demonstrated a reduction of metastability and functional network repertoire in UWS compared to MCS patients. This was expressed in terms of diminished dwell times and loss of nonstationarity in the default mode network and fronto-parietal subcortical network in UWS compared to MCS patients. We further demonstrated that these findings co-occurred with a loss of dynamic interplay between structural eigenmodes and emerging time-resolved functional connectivity in UWS. These results are, amongst others, in support of the global neuronal workspace theory and the mesocircuit hypothesis, underpinning the role of time-resolved thalamo-cortical connections and metastability in the recovery of consciousness.

## I. Introduction

Diagnosis of the level of consciousness after coma due to severe brain injury is a well-known dilemma in the field of neurology and intensive care medicine. Coma after cardiac arrest or after traumatic brain injury (TBI) may result in sustained altered states of consciousness. These patients with disorders of consciousness (DOC), irrespective of the aetiology, can be grouped into the unresponsive wakefulness syndrome (UWS) (1), characterized by the presence of eye-opening and reflexive behaviours, and the minimally conscious state (MCS), characterized by consistent but fluctuant wilful conscious behaviours, such as command following or visual pursuit (2, 3). Recovery of consciousness is argued to emerge conjointly with restoration of resting-state functional brain networks (4), which refers to patterns of neuronal interactions inferred by indirect (e.g., functional magnetic resonance imaging - fMRI) or direct (e.g., electro- and magneto-encephalography (EEG/MEG)) measurements. Analysis of these resting-state networks could potentially help in the diagnosis of patients with DOC and provide insight into the mechanisms that results in absence of recovery of consciousness in UWS.

Various resting-state networks that play an important role in the recovery of consciousness have been identified, among which the default mode network (DMN), fronto-parietal network (FPN) and the salience network are the most important (5, 6). Recovery of the DMN in combination with recovery of the auditory network could for instance discriminate between MCS and UWS with a very high accuracy (~85%) (7). The mechanism of resting-state network restoration in DOC is yet unknown, however, thalamic activity and especially thalamocortical connectivity may be a driving force behind restorations of cortical network function that sustains conscious states (8, 9). Previous work on resting-state networks in DOC have mainly focused on the “*static*” picture of functional connectivity (4, 6, 10, 11), i.e., connections are assessed over the entire duration of the (fMRI) recording and fluctuations in connectivity over time are ignored. However, the underlying dynamics of connectivity seem relevant for consciousness (12, 13) and a static description may therefore be inadequate to provide mechanistic insight into failure of recovery of consciousness in DOC (14).

The analysis of dynamic or time-resolved functional connectivity, as well as the relationship between the underlying anatomical connections and emergent time-resolved functional connectivity (15, 16), may be clinically relevant in patients with DOC. Previous studies have already explored the role of time-resolved functional connectivity in DOC (17–19). A recent study demonstrated that network states with long distance connections occurred less frequently over time in MCS compared to UWS patients (14), emphasizing disintegration of interactions across the cortex in unconscious states. However, network states reminiscent of the well-known resting-state networks were not retrieved. Cao and colleagues used two methods to extract time-varying networks, i.e. independent component analysis and hidden Markov modelling, and revealed clinically relevant differences in network state durations between patients with DOC patients and healthy subjects (20), while lacking comparative analysis between patients in MCS and in UWS. In another fMRI study, the authors focused on the posterior cingulate area and the DMN using a spatiotemporal point process analysis and demonstrated decreased occurrence of DMN-like patterns in UWS. Dynamic connectivity analysis has recently also been applied to EEG data, revealing loss of network integration and increased network segregation in DOC patients (21). Despite the importance of the previously published work, the role of the well-known resting-state networks and especially thalamo-cortical functional connections (22) within the context of time-resolved connectivity and DOC has so far not been fully explored, partly potentially due to the fact that previous work has been mostly hypothesis driven rather than data driven.

Another important aspect in the context of the emergence or restoration of resting-state networks is the underlying structural network, as anatomical connectivity patterns influence the repertoire of possible functional network states (23). It is widely assumed that switching between functional network states is achieved by so-called metastability in the brain (24), i.e., winnerless competitive dynamics. A promising and robust approach to analyse the relationship between structural and functional network states is the so-called eigenmode approach (25–27). With this approach, spatial harmonic components or eigenmodes are extracted from the anatomical network. These eigenmodes can be considered as patterns of ‘hidden connectivity’, which allow for prediction of the well-known resting-state networks (28). It can be hypothesized that switching between functional network states, as can be observed in the metastable brain, is accompanied by fluctuations in the expression of eigenmodes (29); therefore, a potential loss of metastability in DOC could co-occur with loss of modulations in eigenmode expression (12).

In this context, the aim of the current study was fourfold. First, we first tested whether loss of metastability and resting-state network activity, derived from time-resolved estimates of functional connectivity, could differentiate between MCS and UWS, with a potential extraction of a spatiotemporal thalamocortical network state. Second, we analysed whether time-resolved connectivity could be explained by modulations in expression of eigenmodes in DOC, and third, whether potential differences in eigenmode expression in DOC patients would co-occur with a loss of metastability. Finally, we used a classification procedure to evaluate whether the collection of these network-based measures could predict patients’ diagnosis.

## II. Results

We included 34 healthy control subjects (HC, 39 (mean) ± 14 years (standard deviation), 20 males), 30 MCS (41 ± 13 years, 21 males) and 14 UWS patients (48 ± 16 years, 7 males). There was no difference in the age of patients with MCS and UWS (*p* > 0.05), gender (*p* > 0.05), time since injury (*p* > 0.05) and aetiology (*p* > 0.05). There was also no difference in age (*p* > 0.05) and gender (*p* > 0.05) between HC and DOC patients. Further details about the patient population is described in the methods and supplementary table 1.

### Metastability and time-resolved functional connectivity in patients with DOC

Time-resolved or dynamic connectivity for all subjects was extracted from the phase information of the data. We quantified a proxy measure for metastability defined as the standard deviation of the overall phase behaviour over time (i.e. the Kuramoto order parameter). This was followed by extraction of spatiotemporal patterns using non-negative tensor factorisation (NNTF) from phase connectivity data, corresponding to resting-state networks or network states (see Figure 1). Well-known resting-state networks as well as a residual component were used as initial conditions for the spatial connectivity patterns for all network states to allow for stable convergence of the algorithm (i.e. DMN, FPN, visual network, sensorimotor network, salience network, subcortical network (30)). However, the NNTF algorithm allowed the spatial patterns of these network states to change in order to maximize the explained variance of the data. Temporal statistics from the network states were derived for every network state in terms of excursions from the median (proxy for nonstationarity (31) and state duration (i.e., dwell time).

**Figure 1:**
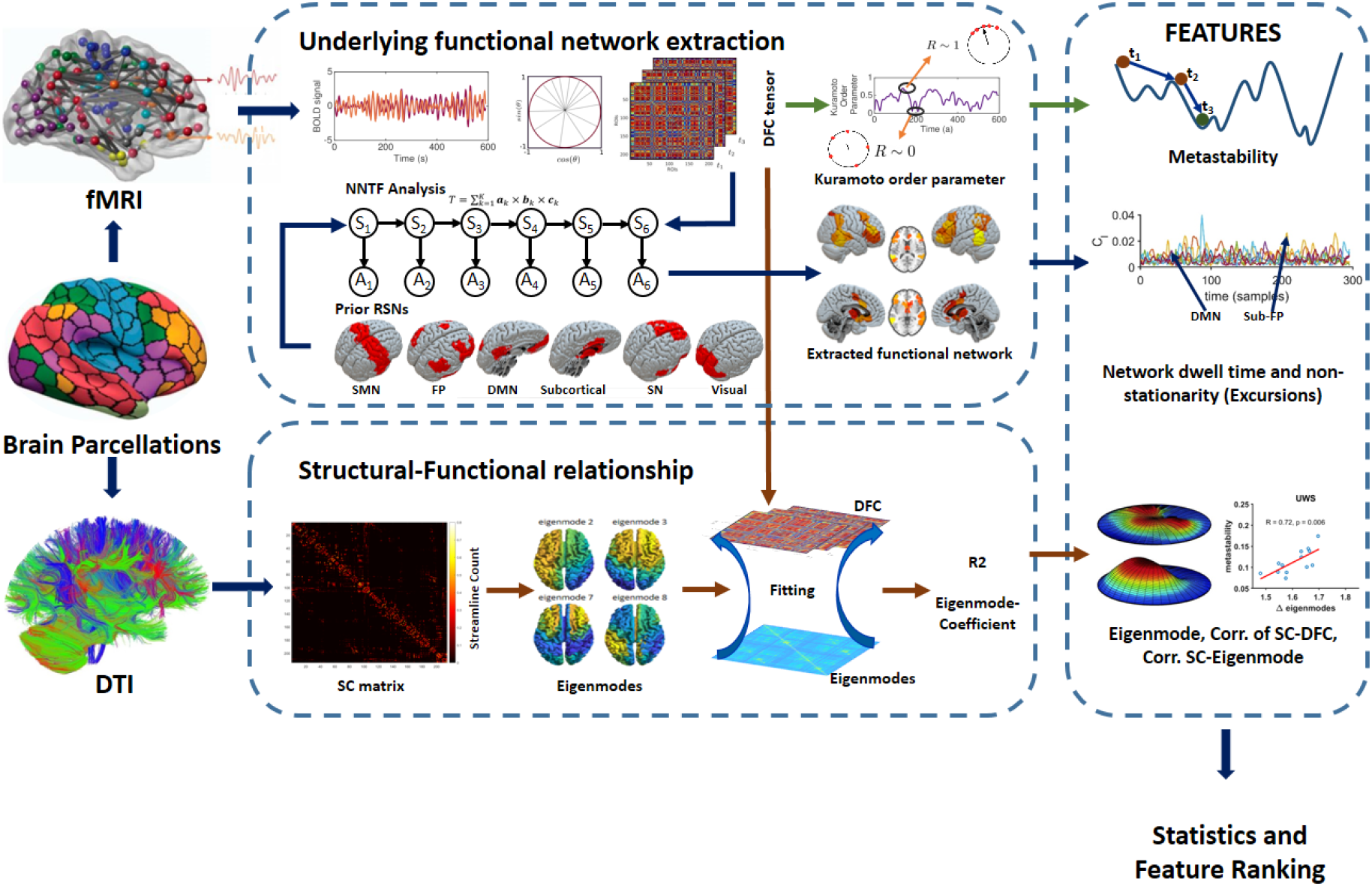
Overview of the analysis pipeline. We used the same Shen parcellation for DWI and fMRI data. Time-resolved functional connectivity was estimated using a metric for phase connectivity. A proxy measure for metastability was derived from the phase information. Time-resolved networks were subsequently extracted from the concatenated data from all subjects using non-negative tensor factorisation (NNTF). Dwell times and nonstationarity (excursions from the median) were retrieved for each spatial pattern of functional connectivity (6 resting-state networks and 1 ‘residual’ network). At the same time, time-resolved connectivity was predicted on a sample by sample basis based on a linear combination of eigenmodes (hidden patterns in the anatomical network). Measures were used for classification and feature ranking.

A reduction of metastability was found in DOC patients compared to HCs (Figure 2A). Lower metastability was observed in UWS patients in comparison to MCS patients (Figure 2A). Reduced metastability is expected to occur with loss of switching between resting-state networks and potentially with dwelling within a more limited subset of resting-state networks in DOC. The output of the NNTF algorithm resulted in spatial topographies of some of the well-known resting-state networks, i.e., the default mode network (DMN), a separate posterior DMN around the precuneus, the visual network, the salience network (SN), the fronto-parietal network (FPN), and a network consisting of fronto-parietal and subcortical regions (FPN-sub) (Figure 2 I-N). Note that these network states were not identical to the initial conditions, e.g. the subcortical network that was provided as initial condition to NNTF was incorporated with the frontoparietal network (Figure 2N) by the NNTF algorithm. At the same time, the sensorimotor network that was provided as initial condition disappeared as state. Excursions from the median were lower for most networks (DMN, visual, Salience, Posterior DMN and FPN-sub) in DOC compared to HC (Figure 2 C-F, H). Significant loss of nonstationarity was also found in UWS compared to MCS for the DMN, FPN, FPN-sub (Figure 2 C,G,H). The NNTF also yielded a residual state, with a lack of spatial structure, accounting for the variance of connectivity data not explained by the resting-state networks. The residual component had longer dwell times for the decreasing levels of consciousness (Figure 2B). In addition, as in line with the metastability results, there were lower dwell times in DOC patients for a specific set of resting-state networks (Salience, Posterior DMN, FPN and FPN-sub), and dwell time was shorter in UWS patients compared to MCS patients only in the FPN-sub network (see results in Figure S1).

**Figure 2:**
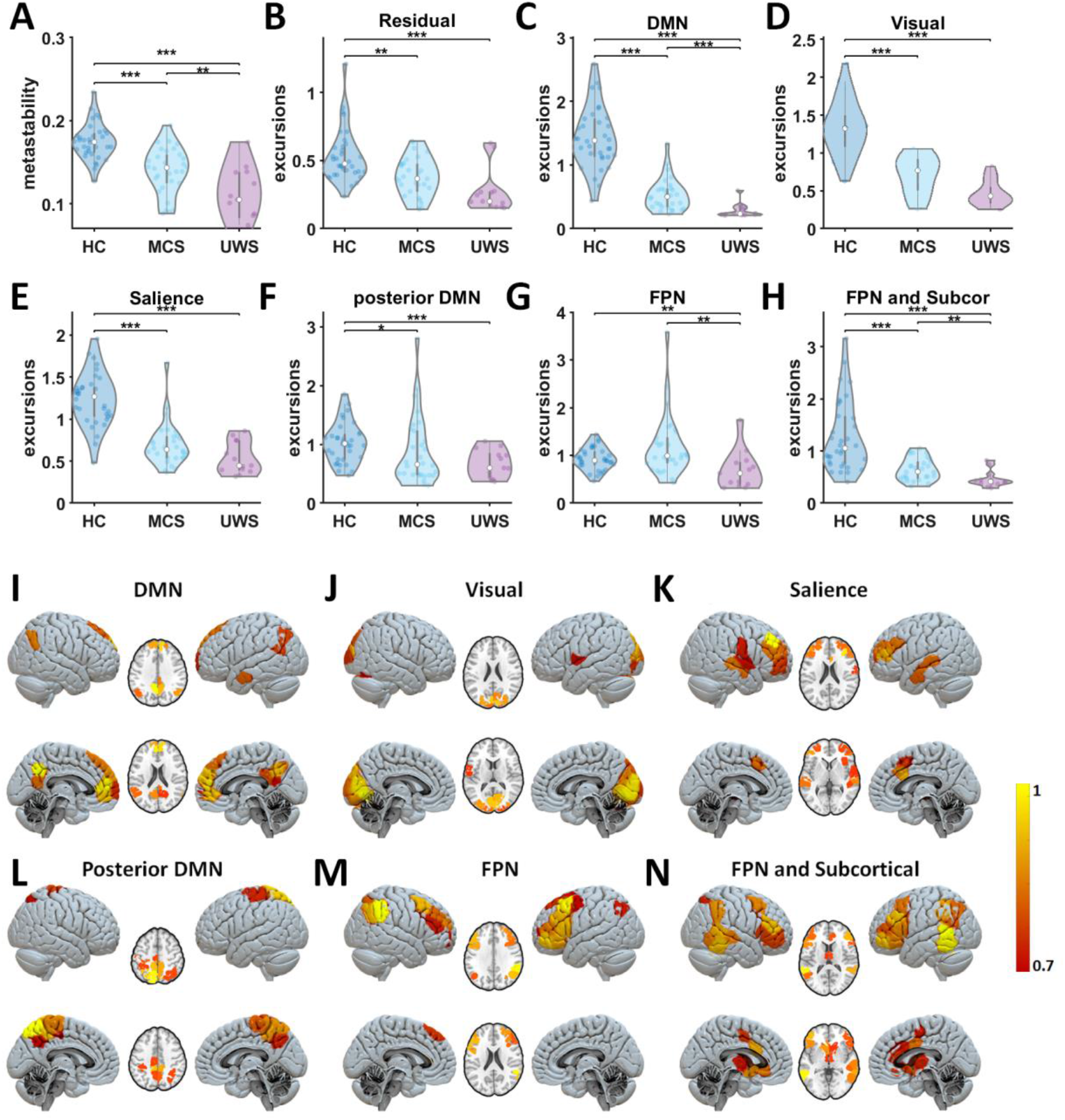
Metastability and time-resolved functional networks in DOC. Panel A displays metastability for all groups: healthy controls (HC), minimally conscious state (MCS), unresponsive wakefulness state (UWS). Panels B-H display the distributions of nonstationarity (excursions from the median) for the residuals and time-resolved networks. Panels I-N show the spatial patterns of time-resolved output networks. Abbreviations: default mode network (DMN), frontoparietal network (FPN), **, and *** denote p < 0.01 and p <0.001 respectively. The colourbar indicates the strength of that specific area to the overall spatial pattern.

### Relationship between structural eigenmodes and time-resolved functional connectivity in DOC

We next analysed how disruption in time-resolved functional connectivity in DOC was related to the underlying structural network. In order to put our findings into context, we first analysed the relationship between static functional networks and structural networks, using the Pearson correlation between static functional connectivity and structural connectivity for the different groups (Figure 3A). These results show that functional connectivity in DOC patients show more correspondence with the underlying structural connectivity as compared to HCs, as the relationship between structural and functional connectivity was stronger for decreasing levels of consciousness (Figure 3A).

**Figure 3:**
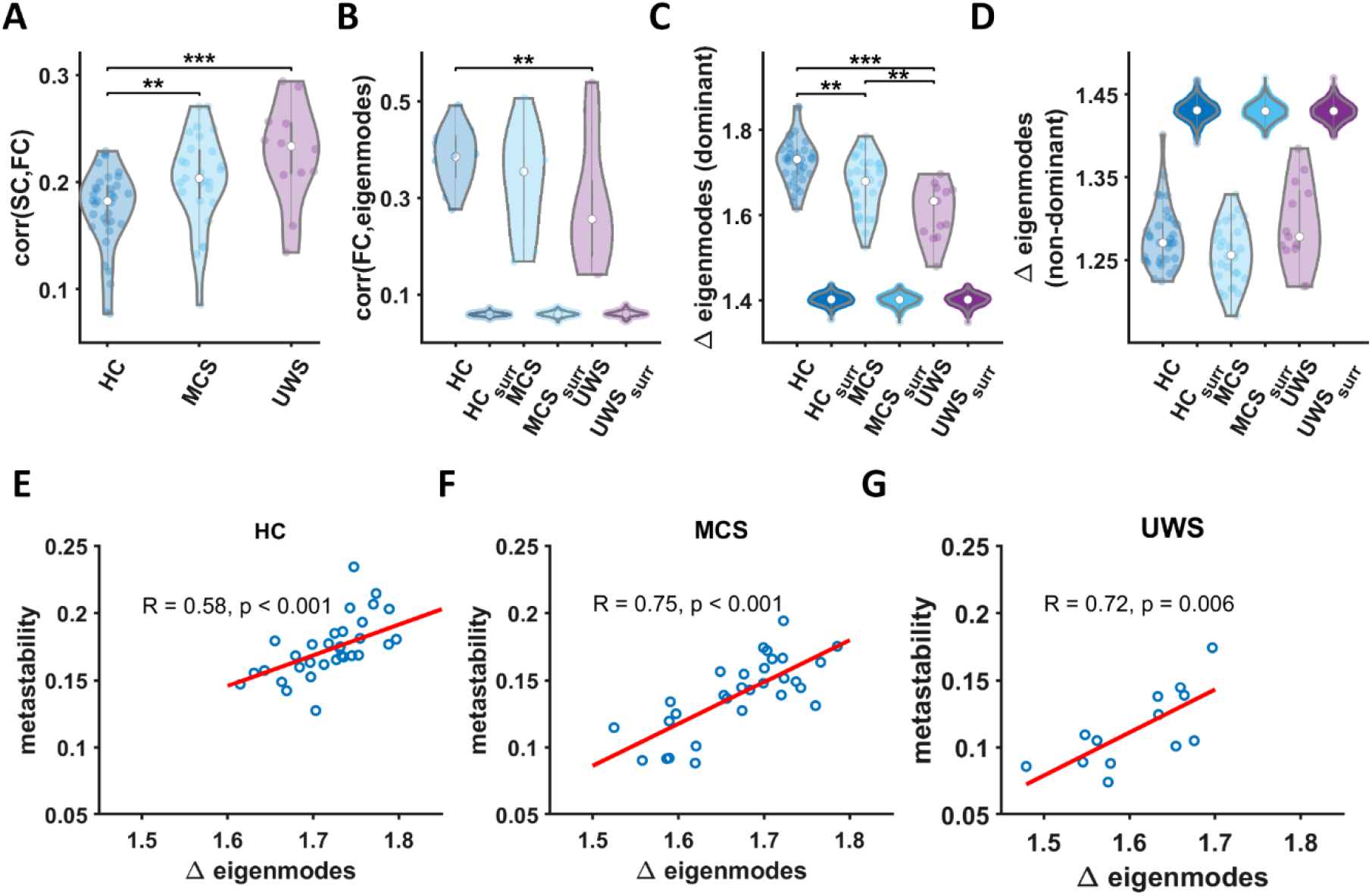
Relationship between time-resolved connectivity and eigenmodes. Panel A shows the prediction of static functional connectivity based on structural connectivity for all three groups (HC, MCS and UWS) in terms of the Pearson correlation coefficient. Panel B shows the prediction of time-resolved functional connectivity based on eigenmodes. These distributions of eigenmode predictions are accompanied with predictions based on surrogate data. We further illustrate the level of fluctuations in eigenmode expression for all three groups (HC, MCS, UWS) for dominant (reflecting network integration, Panel C) and non-dominant eigenmodes (reflecting increasing network segregation, Panel D) accompanied with results for surrogate data, ** and *** denote p < 0.01 and p <0.001 respectively. Panels E-G show that metastability is strongly correlated to modulations in eigenmode expression within every group.

We next obtained the eigenmodes from the structural connectivity by extracting the eigenvectors of the graph Laplacian. These eigenmodes can be regarded as distinct spatial harmonics within the structural connectivity, where the first eigenmodes correspond to patterns with low spatial frequency and subsequent eigenmodes contain patterns with increasingly higher spatial frequencies. Given their spatial configuration, consecutive eigenmodes can be associated with increasing levels of segregation while the first eigenmodes can be linked with network integration. For every time point we predicted the extent to which phase connectivity could be explained by a weighted combination of the eigenmodes (27). Since phase connectivity can evolve over time, the weighting coefficients for the eigenmodes can modulate as well, resulting in fluctuations in the strength of the expressions of eigenmodes over time. For every eigenmode, we could then quantify the modulation strength (i.e. how much the eigenmode-expression varied over time). In addition to the weighting coefficients, we also obtain the goodness-of-fit for the predictions of time-resolved functional connectivity.

The goodness-of-fit for the eigenmode predictions is displayed in Figure 3B, where we show the average correlation between eigenmode predicted FC and empirical FC for the three groups. Results show better predictions for HC and MCS compared to predictions for static FC (median and interquartile range of correlations HC static 0.18 ± 0.04, HC eigenmode 0.39 ± 0.09, Z = −7.1, *p* < 0.001, MCS static 0.2 ± 0.05, MCS eigenmode 0.35 ± 0.18, Z = −4.8, *p* < 0.001). In order to test whether these eigenmode predictions of time-varying connectivity could have been obtained by chance, we redid our analysis using surrogate BOLD data (see method section “Analysis steps”). Results showed that eigenmode predictions for time-resolved connectivity from surrogate data performed significantly worse compared to genuine empirical data (for all comparisons with surrogate data *p* < 0.001; Figure 3B). We did not test whether contribution of individual eigenmodes differed between groups as this would come with a serious multiple comparisons problem.

Instead, since structural connectivity appeared to correlate stronger with static FC in DOC compared to HC, we expected that eigenmode coefficients in DOC patients would hardly change over time, underlining the observation of a ‘fixed’ structural-functional network relationship in DOC patients. To analyse this lack of change in the structural-functional network relationship over time in DOC patients, we quantified the modulation strength of the weighting coefficients over time (see methods “*Analysis steps*”). We performed this analysis separately for the dominant (1^st^ to 107^th^ eigenmode, first half) and non-dominant eigenmodes (108^th^ to 214^th^ eigenmode, second half). Results for the dominant eigenmodes show a clear reduction in modulation of the eigenmode weighting in DOC patients compared to HCs (Figure 3C), with also a significantly lower modulation of eigenmode expression in UWS compared to MCS patients. This result could not be explained by chance, since the same results could not be obtained from surrogate data (Figure 3C). Note that no between group difference for non-dominant eigenmodes was obtained (Figure 3D).

We have so far shown a reduction in modulation strength of eigenmode expressions in DOC patients compared to HC subjects, as well as a loss of metastability in DOC patients and dwelling of the brain in fewer network states in DOC patients. This poses the question whether these two observations are related. In Figure 3EFG we show that metastability is strongly correlated to modulations in eigenmode expression within every group. This underscores the notion that loss of dynamic modulations in functional network patterns due to a loss of metastability could indeed be related to a reduced modulation of eigenmode expression.

### Classification of DOC patients using measures of dynamic functional connectivity and structure-function relationships

To translate the structural and functional dynamic properties of the brain to clinical practice, we used a classification approach for functional and structural properties using two class support vector machine (SVM) classifiers. Only functional and structural properties that showed group differences were used as features. We used three different classification approaches of SVM (i.e., leave one out cross validation (LOOCV), k-fold cross validation, and splitting the data into 60-40% training and testing data respectively). The LOOCV showed better performance in terms of classification between groups for the selected features, compared to the other approaches (see Table 1). Using LOOCV, UWS versus MCS classification accuracy was 79.1%, with a sensitivity of 83.3% and specificity of 69.2%. When we compared healthy controls with the DOC patient group, the classification performance was very high. For healthy controls versus UWS, we found a classification accuracy of 95.8%, with a sensitivity of 97.1% and a specificity of 92.3%. The healthy controls versus MCS classification accuracy was 95.3%, with a sensitivity of 94.1% and specificity of 96.7% (Table 1). We also used the surrogate data to assess whether such a classification accuracy could be obtained by chance. We found that the classification algorithm assorted all subjects into one group, with an accuracy equal to chance level, a sensitivity of 100% and specificity of 0% (Table 1). These results indicate that the classification performance of functional and structural features is beyond chance level.

**Table 1.** Classification accuracy, sensitivity and specificity (mean and standard deviation) for MCS vs UWS, HC vs UWS and HC vs MCS using three different SVM-based classification approaches. Performance of the classification using the real experimental data is presented alongside the performance of classicisation based on surrogate data, to define change level.

| SVM classification method | Group Comparison | Accuracy |  | Sensitivity |  | Specificity |  |
| --- | --- | --- | --- | --- | --- | --- | --- |
|  |  | Experimental data | Surrogate data | Experimental data | Surrogate data | Experimental data | Surrogate data |
| LOOCV (Leave-one-out cross-validation) | MCS vs UWS | 79.1% | 66% | 83.3% | 100% | 69.2% | 0% |
|  | HC vs UWS | 95.8% | 72% | 97.1% | 100% | 92.3% | 0% |
|  | HC vs MCS | 95.3% | 53% | 94.1% | 100% | 96.7% | 0% |
| 10-fold cross-validation | MCS vs UWS | 76.1±0.03 % | 66±0% | 80.0±0.04 % | 100±0% | 67.1±0.04 % | 0±0% |
|  | HC vs UWS | 95.5±0.01 % | 72±0% | 96.6±0.01 % | 100±0% | 92.8±0.04 % | 0±0% |
|  | HC vs MCS | 93.8±0.02 % | 53±0% | 93.8±0.01 % | 100±0% | 93.7±0.03 % | 0±0% |
| 60-40 data splitting | MCS vs UWS | 73.5±0.12 % | 66±0% | 78.4±0.14 % | 100±0% | 64.5±0.11 % | 0±0% |
|  | HC vs UWS | 95.5±0.05 % | 72±0% | 96.3±0.07 % | 100±0% | 93.4±0.08 % | 0±0% |
|  | HC vs MCS | 93.6±0.05 % | 53±0% | 93.3±0.06 % | 100±0% | 94.2±0.08 % | 0±0% |

To further understand which features were most discriminating between UWS and MCS, we used a feature ranking based on diagonal adaptation of neighbourhood component analysis (NCA). Results showed that the most important features were nonstationarity in the DMN (feature weight (FW)=2.22), Salience network (FW=1.03), FPN (FW=0.66), visual network (FW=0.18), FPN-sub network (FW=0.1). Remaining features had low feature weights (<0.01) (Figure 4). Interestingly we found that purely structural features had very low weights, indicating that purely structural properties contribute very little beyond functional features to the classification between UWS versus MCS, as well as between healthy controls versus DOC patients.

**Figure 4:**
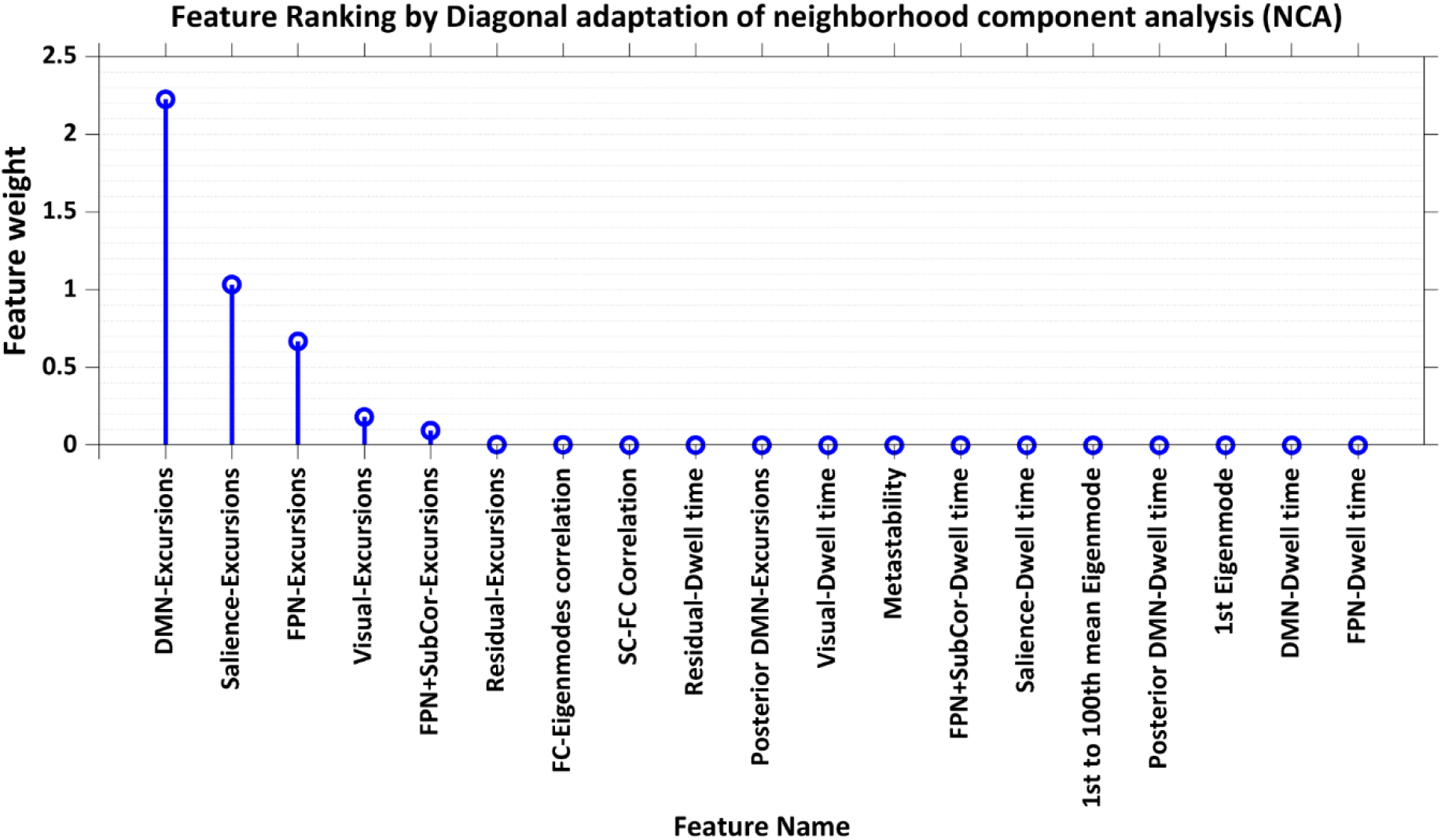
Feature ranking. We illustrate the feature weights for classification based on diagonal adaptation of neighborhood component analysis (NCA).

## III. Discussion

Differentiation between MCS and UWS is key for adequate diagnosis and prognosis in DOC patients as this is connected to medical-ethical end of life decisions. Use of imaging characteristics allows testing of hypotheses on causes for delayed, or failure of, recovery of consciousness. Here, we used state-of-the art techniques to quantify time-varying functional connectivity, metastability and the relationship between the underlying anatomical network and time-resolved functional connections. We demonstrated that these advanced techniques were sensitive to detect clinically relevant differences for the diagnosis of MCS and UWS patients. More specifically, we first demonstrated that UWS patients show reduced metastability, and spend less time in states outside the natural equilibrium state that would favor cerebral processing in a cooperative and coordinated manner to support consciousness. This is accompanied by shorter state durations that the brain spends in the frontoparietal-subcortical configuration in UWS. A loss of nonstationarity was observed in several resting-state networks (i.e., DMN, frontoparietal and frontoparietal-subcortical) in UWS compared to MCS patients. We furthermore showed that functional brain networks are more ‘fixed’ to the underlying anatomical connections and are less subject to spatial reconfigurations over time in UWS compared to MCS patients. The extent to which these spatial reconfigurations occurred (i.e., expressed as modulations in eigenmode-expression) correlated strongly to metastability. Lastly, classification analysis showed that out of all results, nonstationarity in the DMN, salience network, frontoparietal network, visual network and in the frontoparietal-subcortical network were features that were most discriminating between MCS and UWS.

Our results are in agreement with several hypothesis and theories for the emergence of consciousness, of which most share the importance of thalamocortical connectivity for consciousness (32–34). The mesocircuit hypothesis states that deafferentation between the frontal cortex and subcortical regions is crucial in explaining failure of recovery of consciousness (32). One of the most novel findings in the current work is the generation of the frontoparietal-subcortical network. Although subcortical connections were, among others, used as initial conditions for the decomposition of the time-varying functional connectivity patterns into resting-state networks, incorporation with fronto-parietal connections emerged from the data-driven NNTF algorithm. Another observation confirms that this NNTF approach was extracting DOC-relevant networks, namely that the sensorimotor network disappeared after optimization of spatial network patterns. This latter result is in line with the fact that somatosensory cortices are not directly involved in the emergence of consciousness, based on current theories (35). In addition, we found that the frontoparietal-subcortical network showed shorter dwell times in DOC patients compared to HC subjects, with even shorter state durations in UWS compared to MCS patients. Finally, this network also demonstrated a loss of nonstationarity in UWS compared to MCS patients. However, it should be noted that the frontoparietal-subcortical network was not the only network with loss of time-resolved network characteristics; other resting-state networks also showed loss of nonstationarity, such as the DMN and frontoparietal network. Yet a combination of shorter dwell times and loss of nonstationarity was only found for the frontoparietal-subcortical network. The mesocircuit hypothesis suggests that lack of excitation of the inhibition of the thalamus induces a reduction of thalamo-cortical connectivity, which in turn, causes a reduction of the activity in the whole frontoparietal network. It may be tempting to interpret that the frontoparietal-subcortical network may play a crucial role in orchestrating global network interactions and dwell times. Hence, this sub-network may be instrumental for the observed loss of nonstationarity in the other sub-networks. Although the importance of functional connections between the thalamus and frontal cortex has been emphasized by the mesocircuit hypothesis, and shown to relate to consciousness in hypothesis-driven functional (e.g., (8, 22, 36)) and structural (e.g., (37, 38)) neuroimaging studies, this is the first demonstration of the ability of a (semi)data-driven approach to identify this sub-network in the context of time-resolved functional connectivity. Most previous data-driven approaches have been unable to extract such a network (14, 20, 21).

Our findings also support the global neuronal workspace (GNW) theory (33), which emphasizes the importance of long-distance and recurrent functional connections, large-scale reverberant networks and metastable brain states in the emergence and recovery of consciousness (39). So far, the importance of metastability has mainly been addressed in the context of recovery of consciousness from anesthesia (40). Here, we underscore this finding and demonstrate that a reduction of metastability can even differentiate between UWS and MCS patients. We further show that in DOC patients, the brain is dwelling shorter in relevant and important network states, indicating that a more limited repertoire of functional network states in DOC patients. There is still some dwelling time in the DMN, salience network, visual network and in the unstructured residual state, but shorter dwell time in especially the frontoparietal and frontoparietal-subcortical network.

Diagnosis and prognosis of patients in DOC is challenging and merely relying on clinical measures may be unreliable (41, 42). Specialistic imaging techniques such as positron emission tomography (PET) have shown their added-value to complement the clinical diagnosis of DOC patients (4). Especially the lack of activation of a frontoparietal-subcortical network in UWS has been postulated by several hypotheses on DOC (4). Here, we have used a non-invasive imaging protocol and demonstrated the role of time-resolved functional connectivity and disrupted structural-functional network coupling to differentiate between MCS and UWS patients. Our findings allow differentiation between MCS and UWS with about 80% accuracy, commensurate with previous work (7, 37, 43, 44). A high sensitivity (83%, i.e., true positive for the presence of consciousness) and slightly lower specificity (69%, i.e., true negative predicting the absence of consciousness) was obtained. This might reflect the finding that behavioural assessment might underestimate the presence of consciousness in up to 2/3 of the UWS patients (45). Since our sample size was limited with only 14 UWS patients, we did not use a separate validation dataset to verify our classification results. Instead, we applied a few different approaches to verify our results and we used surrogate data to find out whether our classification could have been obtained by chance. This was not the case. Diagnosis in itself was not the sole goal of the classification analysis, but the adopted approach also aided to elucidate mechanisms that would lead to (failure of) recovery of consciousness in DOC leveraging the data-driven obtained features. Loss of nonstationarity in the DMN, salience network and frontoparietal turned out to be important discriminating features to tell apart MCS and UWS.

Our observation of functional connectivity dynamics that are more restricted to the structural connectivity has been observed in pharmacological and pathological loss of consciousness (see also (14, 46)), however, here we show that this co-occurred with a reduction of metastability. Previous work has demonstrated that the underlying anatomical connectivity forms a constraint for functional connectivity and also shapes the repertoire of possible functional network states (24). The underlying anatomical connectivity contains so-called inherent ‘hidden patterns’ or eigenmodes with different spatial structures. In a dynamical system such as the brain, these eigenmodes, or a combination of eigenmodes, can sequentially be activated or deactivated (25, 47), and thereby shape the repertoire of possible functional network states. We stress that this framework does not imply that there is some fixed relationship or coupling between structure and function, but rather that parts of the anatomical network support the (sequential) formation of specific functional sub-networks, and not only at the level of individual nodes (46). Although the mechanism behind the (de)activation of these spatial eigenmodes remains to be investigated, we posit that a potential underlying mechanism for a loss of the functional repertoire in DOC is the inability to sequentially dwell for prolonged times in a different set of eigenmodes. This inability was even more pronounced for UWS than for MCS.

A few methodological aspects in our retrospective study deserve further discussion. First of all, we did not analyse the contributions of individual eigenmodes for two reasons: i) although earlier studies have demonstrated that a limited set of eigenmodes could already explain observed functional connectivity pattern (25), in our case, assessing group differences between MCS and UWS on the basis of individual eigenmodes would impose a multiple comparisons problem; ii) our analytical approach is based on the assumption that all eigenmodes are necessary in the mapping to functional connectivity instead of a statistical selection of eigenmodes. Second, using fMRI to look at dynamic FC, and particular phase-based FC, may not be optimal. High temporal resolution of EEG/MEG may be able to provide even more reliable estimates of dynamic FC (even of subcortical structures (48) and on the relation between SC and dynamic FC (47). Last, concatenated data from all groups were fed into the NNTF analysis, instead of per group. This assumes that spatial network structure is similar across groups. Even though this is not necessarily the case, the amount of data to allow for stable NNTF results for individual groups was limited, especially for the UWS group. Future (multicentric) studies with more patients should verify whether the decomposition of the dynamic functional connectivity patterns into the observed sub-networks holds for the separate groups. Our approach to concatenate data from all groups made group comparison much easier though, as there was now no need to ‘match’ potentially slightly different networks from the different groups. This also allowed us to focus on networks that were important in DOC.

Taken together, we have demonstrated that a (semi) data-driven approach has extracted clinically meaningful time-resolved functional brain networks. This unique network-based spatiotemporal characterization accounts for structure-function coupling (i.e., eigenmodes), and shows a relationship with brain stability. The measures that differed between UWS and MCS patients most, were the dominant eigenmodes (reflecting structure-function coupling) and time-resolved functional connectivity in the default mode network, frontoparietal network and the subcortical-frontoparietal network. Interestingly, the latter network was generated by the (semi)data-driven approach to better fit the data, and was to sole network to show shorter dwell times in UWS than MCS patients. This suggests that the subcortical-frontoparietal network might play a pivotal role for supporting conscious network interactions, as is in line with several theorethical and hypothesis-driven studies. Future work will be required to assess to what extent these advanced aspects of connectivity can serve as biomarkers to aid diagnosis and prognosis in DOC.

## V. Methods

### Participants

Forty-four adult DOC patients, of whom 30 in Minimally Conscious State (MCS) (11 females, age range 24-83 years; mean age ± SD, 45 ± 16 years) and 14 with the Unresponsive Wakefulness Syndrome (UWS) (6 females, age range 20-74 years; mean age ± SD, 47 ± 16 years) and thirty-four age and gender matched healthy subjects (HC) (14 females, age range 19-72 years; mean age ± SD, 40 ± 14 years) without premorbid neurological problems were included. The local ethics committee from the University Hospital of Liège (Belgium) approved the study. Written informed consent was obtained from all healthy subjects and the legal representative for DOC patients. The same data was used in (46, 49).

The diagnosis of the DOC patients was confirmed through two gold standard approaches (i.e., (i) behavioural and (ii) fluorodeoxyglucose-positron emission tomography (FDG-PET), excluding patients for whom these two diagnostic approaches disagreed. (i) Patients were behaviourally diagnosed through the best of at least five coma recovery scale revised CRS-R assessments, evaluating auditory, visual, motor, oromotor function, communication and arousal (50). (ii) Behavioural diagnosis was complemented with the visual assessment of preserved brain metabolism in the frontoparietal network using FDG-PET as a neurological proxy for consciousness (41). Patient-specific clinical information is presented in Table 1. We only included patients for whom (1) MRI data were recorded without anaesthesia (2) diagnosis was based on at least 5 repetitions of the CRS-R assessment, (3) diagnosed as UWS or MCS, and (4) the FDG-PET diagnosis was in agreement with the clinical diagnosis. We excluded the patients (1) for whom the patients the structural MRI segmentation was incorrect or (2) if there were excessive head movement artefacts during MR recordings. There were 46 MCS patients in which 16 were discarded due to mismatch of PET and CRS-R diagnosis, 8 for failed segmentation and 4 for head movement artefacts. Amongst the 28 UWS patients, 8 were discarded due to mismatch of the PET and CRS-R diagnosis, 4 for failure of segmentation and 2 for head movement artefacts.

### MRI Data Acquisition

For the DOC dataset, structural (T1 and DWI) and functional MRI data was acquired on a Siemens 3T Trio scanner. 3D T1-weighted MP-RAGE images (120 transversal slices, repetition time = 2300 ms, voxel size = 1.0 x 1.0 x 1.2 mm^3^, flip angle = 9°, field of view = 256 x 256 mm^2^) were acquired prior to the 10 minutes of BOLD fMRI resting-state (i.e. task free) (EPI, gradient echo, volumes = 300, repetition time = 2000 ms, echo time = 30 ms, flip angle = 78°, voxel size = 3 x 3 x 3 mm^3^, field of view = 192 × 192 mm^2^, 32 transversal slices). HC subjects were instructed to keep eyes open and to be in relaxed state during the fMRI data acquisition. Last, diffusion weighted MRI (DWI) was acquired in 64 directions (b-value =1,000 s/mm^2^, voxel size = 1.8 x 1.8 x 3.3 mm^3^, field of view 230 x 230 mm^2^, repetition time 5,700 ms, echo time 87 ms, 45 transverse slices, 128 x 128 voxel matrix) preceded by a single unweighted image (b0).

### Resting -state fMRI preprocessing

Preprocessing was performed as in (46) using MELODIC (Multivariate Exploratory Linear Optimized Decomposition into Independent Components) version 3.14, which is part of FMRIB’s Software Library (FSL, http://fsl.fmrib.ox.ac.uk/fsl). The preprocessing consisted of the following steps: the first five functional images were discarded to reduce scanner inhomogeneity, motion correction was performed using MCFLIRT, non brain tissue was removed using Bet Extraction Tool (BET), temporal band-pass filtering with sigma 100 seconds, spatial smoothing was applied using a 5mm FWHM Gaussian kernel, rigid-body registration was performed, and finally single-session ICA with automatic dimensionality estimation was employed (51). Then, FIX (FMRIB’s ICA-based X-noiseifier) was applied to remove the noise components and the lesion-driven for each subject. Specifically, FSLeyes in Melodic mode was used to manually identify the single-subject independent components (ICs) into “good” for cerebral signal, “bad” for noise or injury-driven artifacts, and “unknown” for ambiguous components. Each component was evaluated based on the spatial map, the time series, and the temporal power spectrum (51). Next, for each subject, FIX was applied with default parameters to remove bad and unknown components. Subsequently, the Shen et al., functional atlas (without cerebellum) was applied for brain parcellation to obtain the BOLD time series of the 214 cortical and subcortical brain areas in each individual’s native EPI space (30, 52). The cleaned functional data were co-registered to the T1-weighted structural image by using FLIRT (53). Then, the T1-weighted image was co-registered to the standard MNI space by using FLIRT (12 DOF), and FNIRT (54). The transformations matrices were inverted and applied to warp the resting-state atlas from MNI space to the single-subject functional data using a nearest-neighbor interpolation method to ensure the preservation of the labels. Finally, the time series for each of the 214 brain areas were extracted using fslmaths and fslmeants.

### Probabilistic Tractography analysis

A whole-brain structural connectivity (SC) matrix was computed for each subject as reported in our previous study (46). Briefly, the b0 image was co-registered to the T1 structural image using FLIRT and the T1 structural image was co-registered to the MNI space using FLIRT and FNIRT(53, 54). The transformation matrices were inverted and applied to warp the resting-state atlas from MNI space to the native diffusion space using a nearest-neighbor interpolation method. Analysis of diffusion images was applied using the FMRIB’s Diffusion Toolbox (FDT) (www.fmrib.ox.ac.uk/fsl). BET was computed, eddy current distortions and head motion were corrected using eddy correct tool (55). Crossing Fibres were modeled by using BEDPOSTX, and the probability of multi-fibre orientations was calculated to enhance the sensitivity of non-dominant fibre populations (56, 57). Then, probabilistic tractography analysis was calculated in native diffusion space using PROBTRACKX to compute the connectivity probability of each brain region to each of the other 213 brain regions. Subsequently, the value of each brain area was divided by its corresponding number of generated tracts to obtain the structural probability matrix. Finally, the SC_pn_ matrix was then symmetrized by computing their transpose SC_np_ and averaging both matrices.

### Metastability and time-resolved functional connectivity

BOLD time series for every ROI *k* were filtered within the narrowband of 0.01-0.9 Hz. We estimated both time-resolved functional connectivity and static functional connectivity. Static functional connectivity (FC_static_) was estimated by the Pearson correlation coefficient between pairwise narrowband time courses. Time-resolved connectivity was estimated from the instantaneous phases. The instantaneous phase of the BOLD signal, *φ_k_*(*t*), was extracted from the analytic signal as obtained from the Hilbert transform. The synchronization between pairs of brain regions was characterised as the difference between their instantaneous phases. At each time point, the phase difference between two regions *j* and *k* was used as estimate for instantaneous phase connectivity Δ*φ_jk_*(*t*) (58). Evaluation of pairwise connectivity at each time point resulted in a functional connectivity tensor (number of ROIs x number of ROIs x time points).

Furthermore, we investigated how the synchronization between different nodes fluctuates across time using the concept of *metastability*. In the current sense, metastability quantifies how variable the states of phase configurations are as a function of time. We quantified metastability (i.e. our proxy measure for metastability) in terms of the standard deviation of the Kuramoto order parameter, 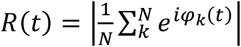, where *N* is the number of ROIs and *i* denotes the imaginary unit (59).

Similar to (60, 61), we extracted time-evolving networks using non-negative tensor factorization (NNTF) (62). We used the N-way toolbox (version 1.8) for MATLAB for this analysis (63). NNTF can be considered as a higher-order principal component analysis. The goal of the approach is to decompose the functional connectivity tensor *FC* into components, such that the approximation of the functional connectivity tensor 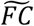 can be written as 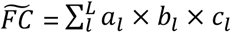. Here, *a_l_* and *b_l_* correspond to vectors decoding spatial information for component *l*, and *c_l_* represents the vector that contains information on temporal fluctuations of component *l*. Note that the outer product *a_l_* × *b_l_* stands for the spatial pattern of functional connectivity of component *l*. The number of components *L* can be estimated from the data using established algorithms (64). However, here, we fixed the number of components based on the number of *a-priori* expected networks that we were interested in. We fed the NNTF algorithm with initial conditions for *a_l_* and *b_l_* based on the spatial components of six expected resting-state networks: salience network, fronto-parietal network, default mode network, subcortical network, sensorimotor network, visual network. We added a residual network to account for the unexplained variance in the functional connectivity tensor. Note that the spatial initial conditions did not indicate that these spatial components were kept fixed during NNTF calculation, but these spatial components were free to be adjusted according to maximization of the explained variance. Data from all subjects and groups were concatenated to allow convergence to stable results.

### Relationship between structural eigenmodes and time-resolved functional connectivity

For every subject, we extracted structural eigenmodes from the graph Laplacian *Q_A_* of the structural connectivity (SC) matrix defined by *Q_A_* = *K_SC_ – SC*, where *K_SC_* refers to the diagonal degree matrix of SC. We further applied symmetric normalization to obtain a normalized Laplacian *Q_sc_*. Subsequently, eigenvectors *u_i_* and eigenvalues (together called eigenmodes) were extracted using diagonalization of *Q_sc_*, resulting in *N* eigenmodes (*N* = number of ROIs). Using a recently introduced approach we mapped functional brain networks at each time point from the structural eigenmodes (27). In other words, we estimated to what extent functional connectivity at each time point *FC*(*t*) could be explained by a linear combination of the eigenmodes

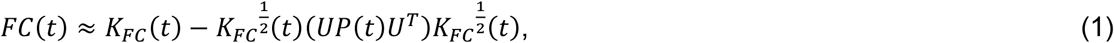

where *K_FC_*(*t*) is the diagonal node strength matrix of the functional connectivity matrix *FC*(*t*) at time *t*. The matrix *P*(*t*) corresponds to the weighting coefficient matrix for the eigenmodes and is obtained after optimisation and equal to

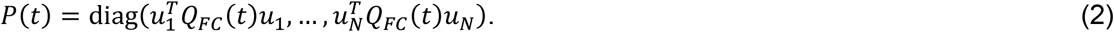

Hence, modulations of eigenmode expressions over time are expressed in *P*(*t*). Here, *Q_FC_*(*t*) is the normalised graph Laplacian of *FC*(*t*) at time *t*, and *u_i_* is the *i-th* eigenvector of *Q_sc_* and the *i-th* column of *U*.

### Analysis steps

1. *Time-resolved functional connectivity*. Individual temporal time courses 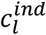 for expression of component *l* for every subject were extracted from *c_l_*. A high value of *c_l_* at a certain time point indicates strong expression of this spatial pattern of functional connectivity (*a_l_* × *b_l_*,) at that time point. At each time point we determined the component with the strongest expression 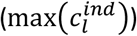, and assumed that connectivity at that point in time was dominated by this state or component. The duration that this component retained the strongest expression was considered as state duration or dwell time (see Figure 1). In addition, we also characterized the amount of nonstationarity in *c_l_* (31, 65), i.e. excursions from the median. The rationale for using this metric is its sensitivity to detect modulations if the underlying system is indeed dynamic (65). Mann-Whitney U tests were used to test, for each network separately, differences in state durations and excursion from the median between groups.
2. *Structural vs functional brain networks*. We first estimated the relationship between static functional connectivity and the structural connectivity itself (without decomposing SC into eigenmodes). This relationship was estimated in terms of a Pearson correlation coefficient between the SC and FC_static_, denoted as corr(FC_static_, SC). We secondly analysed the amount to which time-varying functional networks could be explained by expressions of the structural eigenmodes. We therefore computed the Pearson correlation between the empirical *FC*(*t*) and the eigenmode predicted *FC*(*t*), denoted as corr(FC, eigenmodes). In order to be able to test whether there was a difference in fluctuations of eigenmode expression over time between groups, we quantified the eigenmode modulation strength defined as 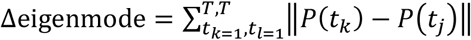, where *t_j_* and *t_k_* correspond to different points in time and *T* to total duration of the recording. As the first eigenmodes can be considered as dominant eigenmodes and more important to shape functional brain networks (25), we evaluated the eigenmode modulation strength for the dominant eigenmodes *i* = {1,…, *N*/2} and non-dominant eigenmodes *i* = {*N*/2,…, *N*} separately. We further computed the correlation between eigenmode modulation strength and metastability in all groups separately in order to test whether modulations in eigenmode expression related to our proxy measure for metastability. Group differences for all metrics was tested using Mann-Whitney U tests. In order to test whether eigenmode predictions of functional connectivity could be obtained by chance, we created surrogate data and redid analysis for the surrogate data. Surrogate data for fMRI BOLD time series were obtained using the circular time shifted method (66). Time-resolved phase connectivity was estimated in the same way as for genuine empirical data. Time-resolved connectivity obtained from surrogate data was subsequently predicted using the eigenmodes. False discovery rate was used to correct for multiple comparisons for analysis steps 1 and 2: number of metrics (excursions * seven networks, dwell time * seven networks, metastability, corr(FC_static_, SC), corr(FC, eigenmodes), non-dominant and dominant eigenmodes)* 3 comparisons + surrogate comparison with genuine data 3 * 3, (corr(FC, eigenmodes), non-dominant and dominant eigenmodes) comparisons = 66 tests) (67).
3. *Classification Algorithm*. For two group classifications (i.e., UWS vs MCS, UWS vs HC, MCS vs HC), a two class “linear SVM” model with 2^nd^ order polynomial kernel was employed. To train the classifier, we used the “fitcsvm” function and to test the classifier performance, we used the “SVMModel.predict” function of MATLAB. As we have a low sample size, we employed three popular algorithms to avoid model or parameter bias. We employed the SVM model with: 1) Leave-one-out cross-validation (LOOCV); 2) 10-fold cross validation and 3) splitting the data in 60-40%. Furthermore, classification performance was verified with real and surrogate data features to get an estimate of the bias in the results obtained with the real data. The discriminative and the predictive capabilities of the classifier were evaluated with measures obtained from receiver operating curves (ROC): accuracy, sensitivity, specificity (68).
4. *Feature Ranking*. To understand which features predominantly contribute to classify the UWS from MCS, and healthy controls from patients, we used the classification-based feature weighting algorithm based on diagonal adaptation of neighbourhood component analysis (NCA). We used the ‘fscnca’ function of MATLAB that learns the feature weights using a diagonal adaptation of NCA and returns the weight for each functional dynamic and structural-functional feature (69).

## References

1. S. Laureys, et al., Unresponsive wakefulness syndrome: a new name for the vegetative state or apallic syndrome. BMC Med. 8, 1–4 (2010).

2. J. T. Giacino, et al., The minimally conscious state: definition and diagnostic criteria. Neurology 58, 349–353 (2002).

3. C. Schnakers, et al., The Nociception Coma Scale: a new tool to assess nociception in disorders of consciousness. Pain 148, 215–219 (2010).

4. B. L. Edlow, J. Claassen, N. D. Schiff, D. M. Greer, Recovery from disorders of consciousness: mechanisms, prognosis and emerging therapies. Nat. Rev. Neurol., 1–22 (2020).

5. E. Amico, et al., Mapping the functional connectome traits of levels of consciousness. Neuroimage 148, 201–211 (2017).

6. L. Heine, et al., Resting state networks and consciousness. Front. Psychol. 3, 295 (2012).

7. A. Demertzi, et al., Intrinsic functional connectivity differentiates minimally conscious from unresponsive patients. Brain 138, 2619–2631 (2015).

8. E. A. Fridman, B. J. Beattie, A. Broft, S. Laureys, N. D. Schiff, Regional cerebral metabolic patterns demonstrate the role of anterior forebrain mesocircuit dysfunction in the severely injured brain. Proc. Natl. Acad. Sci. 111, 6473–6478 (2014).

9. S. Laureys, et al., Restoration of thalamocortical connectivity after recovery from persistent vegetative state. Lancet 355, 1790–1791 (2000).

10. A. Demertzi, A. Soddu, S. Laureys, Consciousness supporting networks. Curr. Opin. Neurobiol. 23, 239–244 (2013).

11. J. T. Giacino, J. J. Fins, S. Laureys, N. D. Schiff, Disorders of consciousness after acquired brain injury: the state of the science. Nat. Rev. Neurol. 10, 99 (2014).

12. P. Barttfeld, et al., Signature of consciousness in the dynamics of resting-state brain activity. Proc. Natl. Acad. Sci. 112, 887–892 (2015).

13. A. I. Luppi, et al., Consciousness-specific dynamic interactions of brain integration and functional diversity. Nat. Commun. 10, 1–12 (2019).

14. A. Demertzi, et al., Human consciousness is supported by dynamic complex patterns of brain signal coordination. Sci. Adv. 5, eaat7603 (2019).

15. L. E. Suárez, R. D. Markello, R. F. Betzel, B. Misic, Linking structure and function in macroscale brain networks. Trends Cogn. Sci. (2020).

16. A. Avena-Koenigsberger, B. Misic, O. Sporns, Communication dynamics in complex brain networks. Nat. Rev. Neurosci. 19, 17 (2018).

17. Y. Sanz Perl, et al., Perturbations in dynamical models of whole-brain activity dissociate between the level and stability of consciousness. PLoS Comput. Biol. 17, e1009139 (2021).

18. D. Golkowski, et al., Dynamic Patterns of Global Brain Communication Differentiate Conscious From Unconscious Patients After Severe Brain Injury. Front. Syst. Neurosci. 15 (2021).

19. S. M. Del Pozo, et al., Unconsciousness reconfigures modular brain network dynamics. Chaos An Interdiscip. J. Nonlinear Sci. 31, 93117 (2021).

20. B. Cao, et al., Abnormal dynamic properties of functional connectivity in disorders of consciousness. NeuroImage Clin. 24, 102071 (2019).

21. J. Rizkallah, et al., Decreased integration of EEG source-space networks in disorders of consciousness. NeuroImage Clin. 23, 101841 (2019).

22. M. M. Monti, et al., Thalamo-frontal connectivity mediates top-down cognitive functions in disorders of consciousness. Neurology 84, 167–173 (2015).

23. G. Deco, et al., Resting-state functional connectivity emerges from structurally and dynamically shaped slow linear fluctuations. J. Neurosci. 33, 11239–11252 (2013).

24. G. Deco, M. L. Kringelbach, Metastability and coherence: extending the communication through coherence hypothesis using a whole-brain computational perspective. Trends Neurosci. 39, 125–135 (2016).

25. S. Atasoy, I. Donnelly, J. Pearson, Human brain networks function in connectome-specific harmonic waves. Nat. Commun. 7(2016).

26. P. A. Robinson, et al., Eigenmodes of brain activity: Neural field theory predictions and comparison with experiment. Neuroimage 142, 79–98 (2016).

27. P. Tewarie, et al., Mapping functional brain networks from the structural connectome: relating the series expansion and eigenmode approaches. Neuroimage 216, 116805 (2020).

28. S. Atasoy, G. Deco, M. L. Kringelbach, J. Pearson, Harmonic brain modes: a unifying framework for linking space and time in brain dynamics. Neurosci. 24, 277–293 (2018).

29. M. G. Preti, D. Van De Ville, Decoupling of brain function from structure reveals regional behavioral specialization in humans. Nat. Commun. 10, 1–7 (2019).

30. E. S. Finn, et al., Functional connectome fingerprinting: identifying individuals using patterns of brain connectivity. Nat. Neurosci. 18, 1664–1671 (2015).

31. A. Zalesky, A. Fornito, L. Cocchi, L. L. Gollo, M. Breakspear, Time-resolved resting-state brain networks. Proc. Natl. Acad. Sci. 111, 10341–10346 (2014).

32. N. D. Schiff, Recovery of consciousness after brain injury: a mesocircuit hypothesis. Trends Neurosci. 33, 1–9 (2010).

33. S. Dehaene, J.-P. Changeux, L. Naccache, The global neuronal workspace model of conscious access: from neuronal architectures to clinical applications. Charact. Conscious. From Cogn. to Clin., 55–84 (2011).

34. H. Blumenfeld, Brain mechanisms of conscious awareness: Detect, pulse, switch, and wave. Neurosci., 10738584211049378 (2021).

35. L. Naccache, Minimally conscious state or cortically mediated state? Brain 141, 949–960 (2018).

36. J. S. Crone, B. J. Bio, P. M. Vespa, E. S. Lutkenhoff, M. M. Monti, Restoration of thalamo-cortical connectivity after brain injury: recovery of consciousness, complex behavior, or passage of time? J. Neurosci. Res. 96, 671–687 (2018).

37. J. Annen, et al., Function–structure connectivity in patients with severe brain injury as measured by MRI-DWI and FDG-PET. Hum. Brain Mapp. 37, 3707–3720 (2016).

38. L. Weng, et al., Abnormal structural connectivity between the basal ganglia, thalamus, and frontal cortex in patients with disorders of consciousness. Cortex 90, 71–87 (2017).

39. G. A. Mashour, P. Roelfsema, J.-P. Changeux, S. Dehaene, Conscious processing and the global neuronal workspace hypothesis. Neuron 105, 776–798 (2020).

40. F. Cavanna, M. G. Vilas, M. Palmucci, E. Tagliazucchi, Dynamic functional connectivity and brain metastability during altered states of consciousness. Neuroimage 180, 383–395 (2018).

41. J. Stender, et al., Diagnostic precision of PET imaging and functional MRI in disorders of consciousness: a clinical validation study. Lancet 384, 514–522 (2014).

42. W. S. van Erp, et al., Unexpected emergence from the vegetative state: delayed discovery rather than late recovery of consciousness. J. Neurol. 266, 3144–3149 (2019).

43. D. A. Engemann, et al., Robust EEG-based cross-site and cross-protocol classification of states of consciousness. Brain 141, 3179–3192 (2018).

44. D. Candia-Rivera, et al., Neural Responses to Heartbeats Detect Residual Signs of Consciousness during Resting State in Postcomatose Patients. J. Neurosci. 41, 5251–5262 (2021).

45. A. Thibaut, et al., Preservation of brain activity in unresponsive patients identifies MCS star. Ann. Neurol. 90, 89–100 (2021).

46. A. López-González, et al., Loss of consciousness reduces the stability of brain hubs and the heterogeneity of brain dynamics. Commun. Biol. 4, 1–15 (2021).

47. P. Tewarie, et al., How do spatially distinct frequency specific MEG networks emerge from one underlying structural connectome? The role of the structural eigenmodes. Neuroimage 186, 211–220 (2019).

48. A. Hillebrand, et al., Detecting epileptiform activity from deeper brain regions in spatially filtered MEG data. Clin. Neurophysiol. Off. J. Int. Fed. Clin. Neurophysiol. 127, 2766–2769 (2016).

49. R. Panda, et al., Posterior integration and thalamo-frontotemporal broadcasting are impaired in disorders of consciousness (2021).

50. J. T. Giacino, K. Kalmar, J. Whyte, The JFK Coma Recovery Scale-Revised: measurement characteristics and diagnostic utility. Arch. Phys. Med. Rehabil. 85, 2020–2029 (2004).

51. L. Griffanti, et al., ICA-based artefact removal and accelerated fMRI acquisition for improved resting state network imaging. Neuroimage 95, 232–247 (2014).

52. X. Shen, F. Tokoglu, X. Papademetris, R. T. Constable, Groupwise whole-brain parcellation from resting-state fMRI data for network node identification. Neuroimage 82, 403–415 (2013).

53. M. Jenkinson, S. Smith, A global optimisation method for robust affine registration of brain images. Med. Image Anal. 5, 143–156 (2001).

54. J. L. R. Andersson, M. Jenkinson, S. Smith, Non-linear registration, aka Spatial normalisation FMRIB technical report TR07JA2. FMRIB Anal. Gr. Univ. Oxford 2, e21 (2007).

55. J. L. R. Andersson, S. N. Sotiropoulos, An integrated approach to correction for off-resonance effects and subject movement in diffusion MR imaging. Neuroimage 125, 1063–1078 (2016).

56. T. E. J. Behrens, et al., Characterization and propagation of uncertainty in diffusion-weighted MR imaging. Magn. Reson. Med. An Off. J. Int. Soc. Magn. Reson. Med. 50, 1077–1088 (2003).

57. T. E. J. Behrens, H. J. Berg, S. Jbabdi, M. F. S. Rushworth, M. W. Woolrich, Probabilistic diffusion tractography with multiple fibre orientations: What can we gain? Neuroimage 34, 144–155 (2007).

58. E. Glerean, J. Salmi, J. M. Lahnakoski, I. P. Jääskeläinen, M. Sams, Functional magnetic resonance imaging phase synchronization as a measure of dynamic functional connectivity. Brain Connect. 2, 91–101 (2012).

59. G. Deco, M. L. Kringelbach, V. K. Jirsa, P. Ritter, The dynamics of resting fluctuations in the brain: metastability and its dynamical cortical core. Sci. Rep. 7, 3095 (2017).

60. A. Ponce-Alvarez, et al., Resting-state temporal synchronization networks emerge from connectivity topology and heterogeneity. PLoS Comput. Biol. 11, e1004100 (2015).

61. P. Tewarie, et al., Tracking dynamic brain networks using high temporal resolution MEG measures of functional connectivity. Neuroimage 200, 38–50 (2019).

62. R. Bro, PARAFAC. Tutorial and applications. Chemom. Intell. Lab. Syst. 38, 149–171 (1997).

63. C. A. Andersson, R. Bro, The N-way toolbox for MATLAB. Chemom. Intell. Lab. Syst. 52, 1–4 (2000).

64. M. E. Timmerman, H. A. L. Kiers, Three-mode principal components analysis: Choosing the numbers of components and sensitivity to local optima. Br. J. Math. Stat. Psychol. 53, 1–16 (2000).

65. R. Hindriks, et al., Can sliding-window correlations reveal dynamic functional connectivity in resting-state fMRI? Neuroimage 127, 242–256 (2016).

66. R. Q. Quiroga, A. Kraskov, T. Kreuz, P. Grassberger, Performance of different synchronization measures in real data: a case study on electroencephalographic signals. Phys. Rev. E 65, 41903 (2002).

67. Y. Benjamini, Y. Hochberg, Controlling the false discovery rate: a practical and powerful approach to multiple testing. J. R. Stat. Soc. Ser. B, 289–300 (1995).

68. R. D. Bharath, et al., Machine learning identifies “rsfMRI epilepsy networks” in temporal lobe epilepsy. Eur. Radiol. 29, 3496–3505 (2019).

69. H. Eryilmaz, et al., Working memory load-dependent changes in cortical network connectivity estimated by machine learning. Neuroimage 217, 116895 (2020).

